# Testing and Correcting for Weak and Pleiotropic Instruments in Two-Sample Multivariable Mendelian Randomisation

**DOI:** 10.1101/2020.04.02.021980

**Authors:** Eleanor Sanderson, Wes Spiller, Jack Bowden

## Abstract

Multivariable Mendelian Randomisation (MVMR) is a form of instrumental variable analysis which estimates the direct effect of multiple exposures on an outcome using genetic variants as instruments. Mendelian Randomisation and MVMR are frequently conducted using two-sample summary data where the association of the genetic variants with the exposures and outcome are obtained from separate samples. If the genetic variants are only weakly associated with the exposures either individually or conditionally, given the other exposures in the model, then standard inverse variance weighting will yield biased estimates for the effect of each exposure. Here we develop a two-sample conditional F-statistic to test whether the genetic variants strongly predict each exposure conditional on the other exposures included in a MVMR model. We show formally that this test is equivalent to the individual level data conditional F-statistic, indicating that conventional rule-of-thumb critical values of F > 10, can be used to test for weak instruments. We then demonstrate how reliable estimates of the causal effect of each exposure on the outcome can be obtained in the presence of weak instruments and pleiotropy, by re-purpousing a commonly used heterogeneity Q-statistic as an estimating equation. Furthermore, the minimised value of this Q-statistic yields an exact test for heterogeneity due to pleiotropy. We illustrate our methods with an application to estimate the causal effect of blood lipid fractions on age related macular degeneration.

## 1 Introduction

Instrumental variables (IV) is a form of regression analysis which estimates the causal effect of an exposure on an outcome in the presence of unobserved confounding. Mendelian randomisation (MR) is a rapidly expanding application of the IV method in the field of epidemiology in which genetic variants are used as instruments. If genetic variants - usually single nucleotide polymorphisms (SNPs) - are available which reliably predict the exposure and are not associated with the outcome through any other pathway, then they are valid IVs. These genetic variants can then be used as instruments to obtain an estimate for the causal effect of a modifiable health exposure on a disease outcome^1,2^. The results of such an analysis can inform the development of public health, or even pharmaceutical, interventions.

Multivariable Mendelian Randomisation (MVMR) is a recently developed extension of MR that can be applied with either individual or summary level data to estimate the effect of multiple, potentially related, exposures on an outcome^3,4^. The three core assumptions that define a set of SNPs, *G*, as valid IV’s for the purpose of an MVMR analysis are;

IV1: *G* must be strongly associated with *each* exposure given the other exposures included in the model;
IV2: *G* is independent of the outcome given *all* of the exposures; and
IV3: *G* is independent of all confounders of *any* of the exposures and the outcome^4^.

These assumptions are illustrated in Fig 1. A violation of IV1 induces ‘weak instrument bias’ in the resulting estimates^5,6^. In a conventional (univariable) MR analysis, the definition of instrument strength is straightforward and unambiguous. Assumption IV1 can be tested with an F-statistic, which tests the association between the SNP and the exposure. When univariable MR analysis based on individual level data from a single sample, if the F-statistic is larger than the rule-of-thumb value of 10 then the SNPs are said to be a ‘strong’ instrument. We can then reject the null hypothesis that the instruments are weak in the sense that the bias of the MR estimate is equal to or greater than 10% of the observational (OLS) association. ^5,6^ In any MVMR analysis it is also necessary that this F-statistic is large for each exposure included, but this is no longer sufficient; the SNPs also need to predict each exposure conditional on the other predicted exposures. This additional condition ensures that there is sufficient variation in association between the SNPs and each exposure, to avoid a problem of weak instrument bias in the MVMR model.

**Figure 1:**
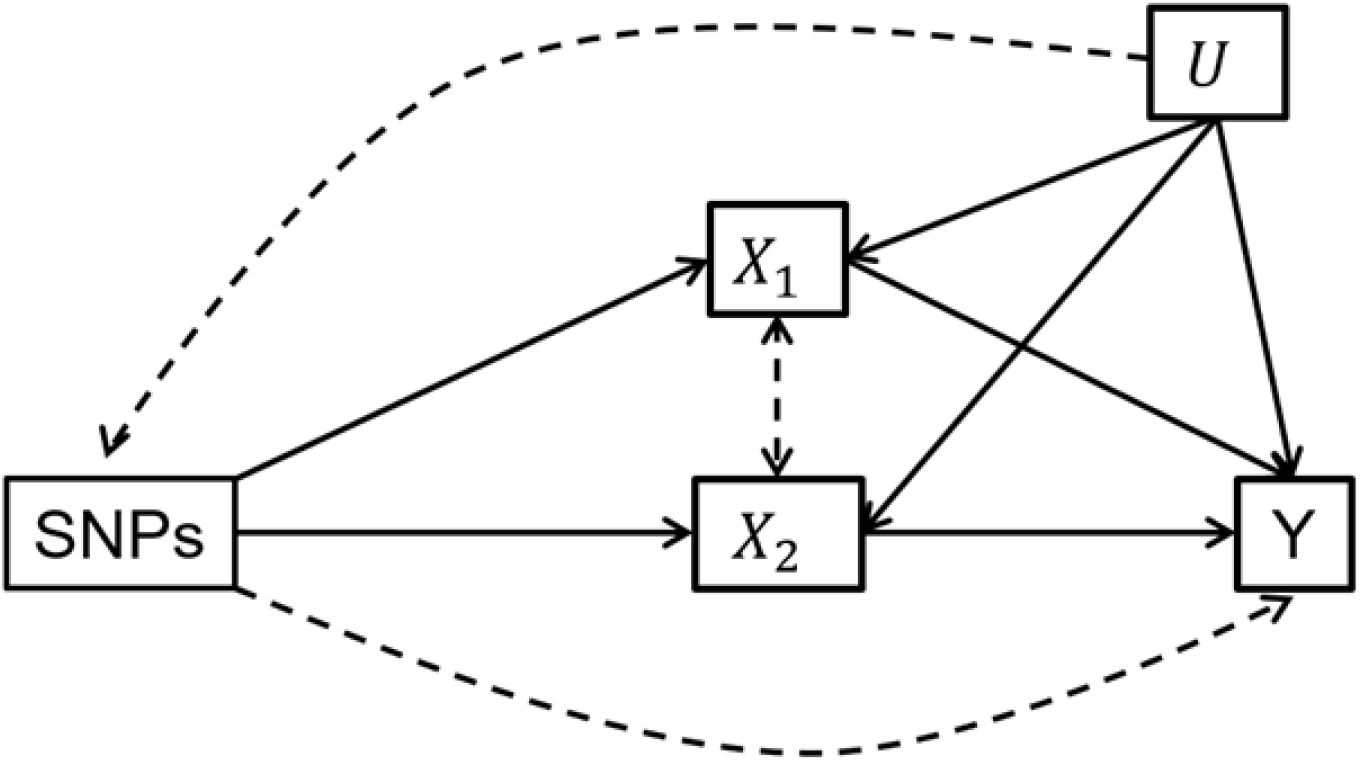
Assumptions for a MVMR analysis DAG illustrating the assumptions required for MVMR. Dashed lines represent associations that must not exist for the SNPs to be valid instruments for the set of exposures

With individual level data, weak instruments can be tested in MVMR using the Sanderson-Windmeijer conditional F-statistic, denoted *F_SW_* ^7,4^. Under weak instruments *F_SW_* has the same distribution as the conventional F-statistic and so can be compared to the same critical values^5,6^. Therefore when testing for weak instruments, verifying that *F_SW_* is greater than the rule-of-thumb of 10 means that we can reject the null hypothesis that the average bias of the MVMR estimates is at least 10% of the bias of the equivalent multivariable OLS estimates.

### Application to two-sample summary data MR

When individual level data on the genetic variants, exposure and outcome are not available two sample MVMR can be conducted using summary data estimates of SNP-exposure and SNP-outcome associations. In two-sample MR, weak instruments bias the causal estimates towards the null rather than the observational association. Sanderson et al (2019) derived a *Q* statistic (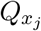) to test for underidentification (i.e. where the SNPs explain none of the variation in the exposures) in two-sample MVMR. We formally show in this paper that a transformation of this statistic has the same distribution as *F_SW_* and therefore can also be compared to the same critical values, or rule-of-thumb of F>10, to test for weak instruments in the two sample setting.

Horizontal pleiotropy is a major threat to the validity of an MR analysis. It occurs when the SNPs have an effect on the outcome (either directly or through another exposure not included in the model) that is not via the exposure of interest, as illustrated by the dashed arrow from *G* to *Y* in Fig 1. This violates assumption IV3 and can lead to biased estimates of the causal effect of each exposure on the outcome from an MR analysis^8^. Although a number of methods currently exist for univariable MR estimation that are robust to pleiotropy under different assumptions^9,10,11,12^ and MVMR can mitigate horizontal pleiotropy via known pleiotropic pathways through the inclusion of multiple exposures, limited methods are available for pleiotropy robust MVMR models^4,13^. Furthermore, in the presence of weak instruments standard tests are increasingly likley to detect pleiotropy when in truth none is present. The major contribution of this paper is to extend weak instrument pleiotropy robust estimation to two sample MVMR with an arbitrary number of exposures. Furthermore, we show that a heterogeneity statistic derived within this estimation procedure provides an exact test for the presence of pleiotropy in the presence of weak instruments. The methods presented here therefore provide the statistical framework for accurate and reliable MVMR model fitting, with potentially large numbers of exposures, in the presence of weak instruments and pleiotropy.

We apply our methods to determine which subset of lipid fractions can be strongly predicted by 150 SNPs associated with at least one of 118 metabolites first presented by Kettunen et al 2016^14^ and estimate the causal effect of those traits on Age related macular degeneration (AMD). The two sample conditional F-statistic calculated for these data helped to highlight that it was not possible to strongly predict multiple lipid fractions from the same subgroup despite each lipid fraction having a moderately high F-statistic. Any analyst naively applying MVMR methods to such data without the correct diagnostic statistics to hand is in danger of generating poor quality and potentially misleading results.

## 2 A Test for Weak Instruments

Let *X* = (*X*_1_*, X*_2_, …, *X_K_*) be a set of *K* exposure variables and let *G* be a set of *L* instruments *G* = (*G*_1_*, G*_2_, …, *G_L_*). Define the *K × L* matrix of associations between each exposure and each instrument as;

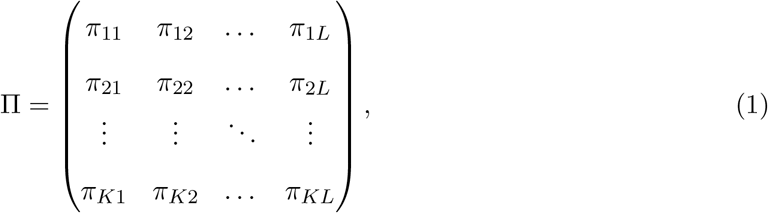

where for example *π*_32_ represents the association between exposure 3 and SNP 2. Without loss of generality, testing whether the instrument set *G* can explain variation in a single exposure, *X*_1_, conditional on all other exposures (*X*_2_, …, *X_K_*) is equivalent to testing whether model (2) below is identified

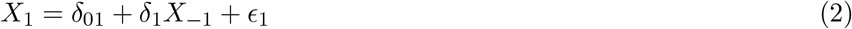

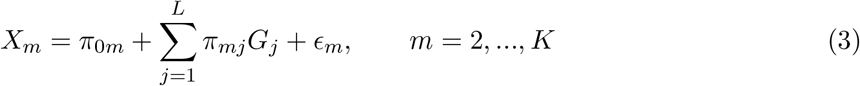

Here: *δ*_01_ and each *π*_0*m*_ are scalar parameters; *δ*_1_ is a *K* − 1 vector of parameters, and *ϵ*_1_ and *ϵ_m_* are random error terms. Collecting *π*_2_, …, *π_K_* into a single (*K* − 1) × *L* matrix, define Π_−1_ as the matrix Π minus its first row. If this model is overidentified then the rank of Π_−1_ is > *K* − (*L* − 1).

In two sample summary data settings we do not directly observe exposures *X*_1_, …, *X_K_*, only estimates for the *K* × *L* SNP-exposure associations that define 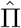. However we can use these association estimates to define an analogous formula to (2)

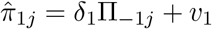

Where Π_−1*j*_ is the *j*th column of Π. The *Q* statistic for exposure 1 based on the summary data estimates can be written as;

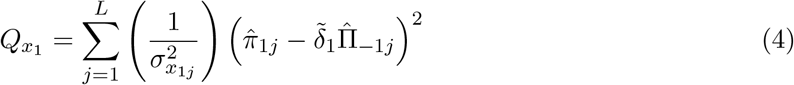

where the variance term 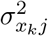 is given by;

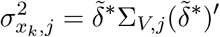

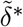 is the K by 1 vector 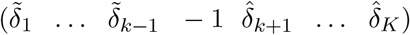, and 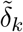 is an efficient estimator for *δ_k_*, for example estimated through an inverse variance weighted least squares regression of 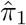 on 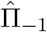. The matrix Σ_*V,j*_ defines the covariance of the estimated effects of snp j on each of the exposures:

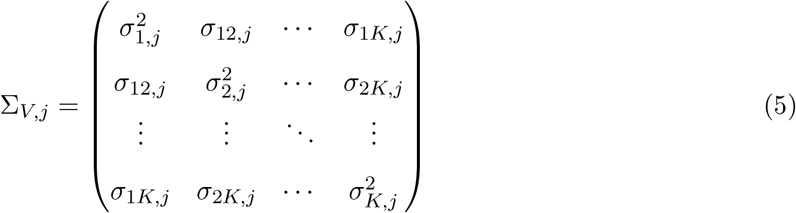

If each 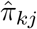 is obtained separately via univariable regressions with an intercept, then the error terms are obtained from the expressions:

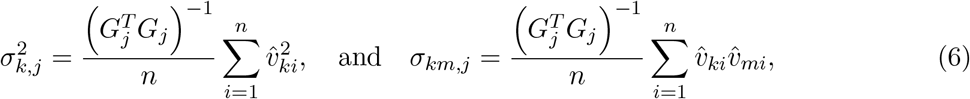

Under the null hypothesis that the instruments do not contain enough information to predict both exposure variables, 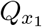 will be asymptotically 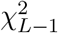 distributed where *L* is the number of SNPs in the estimation. Rejecting the null hypothesis indicates that the SNPs can predict *X*_1_ conditional on *X*_2_. Dividing the Q-statistic described above by the number of instruments, adjusted for the number of exposures, in the model gives a test statistic that is equivalent to the one sample conditional F statistic *F_SW_*.

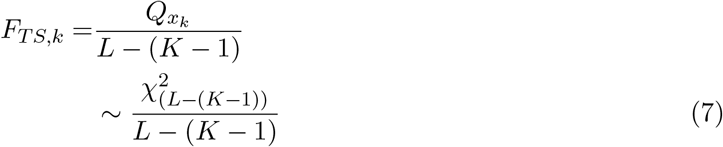

In Supplementary Section S.1 we show that under the assumption that the instruments are uncorrelated the two sample conditional F-statistic *F_TS_* in (7) is equivalent to the one sample conditional F-statistic *F_SW_*. Fig.2 below gives the distribution of the individual conditional F-statistic and the two-sample conditional F-statistic *F_TS_* for models with 25 and 100 SNPs included as instruments. The simulations were generated from a model with two exposures, both of which are strongly individually predicted but jointly weakly predicted by the set of SNPs. That is they had large individual F statistics but small *F_TS_* statistics of 10. The total bias in the two MVMR estimates is therefore approximately 10% of the bias in the observational association. Results are given for one exposure only. This figure supports the formal equivalence result given in Supplementary section S.1.

**Figure 2:**
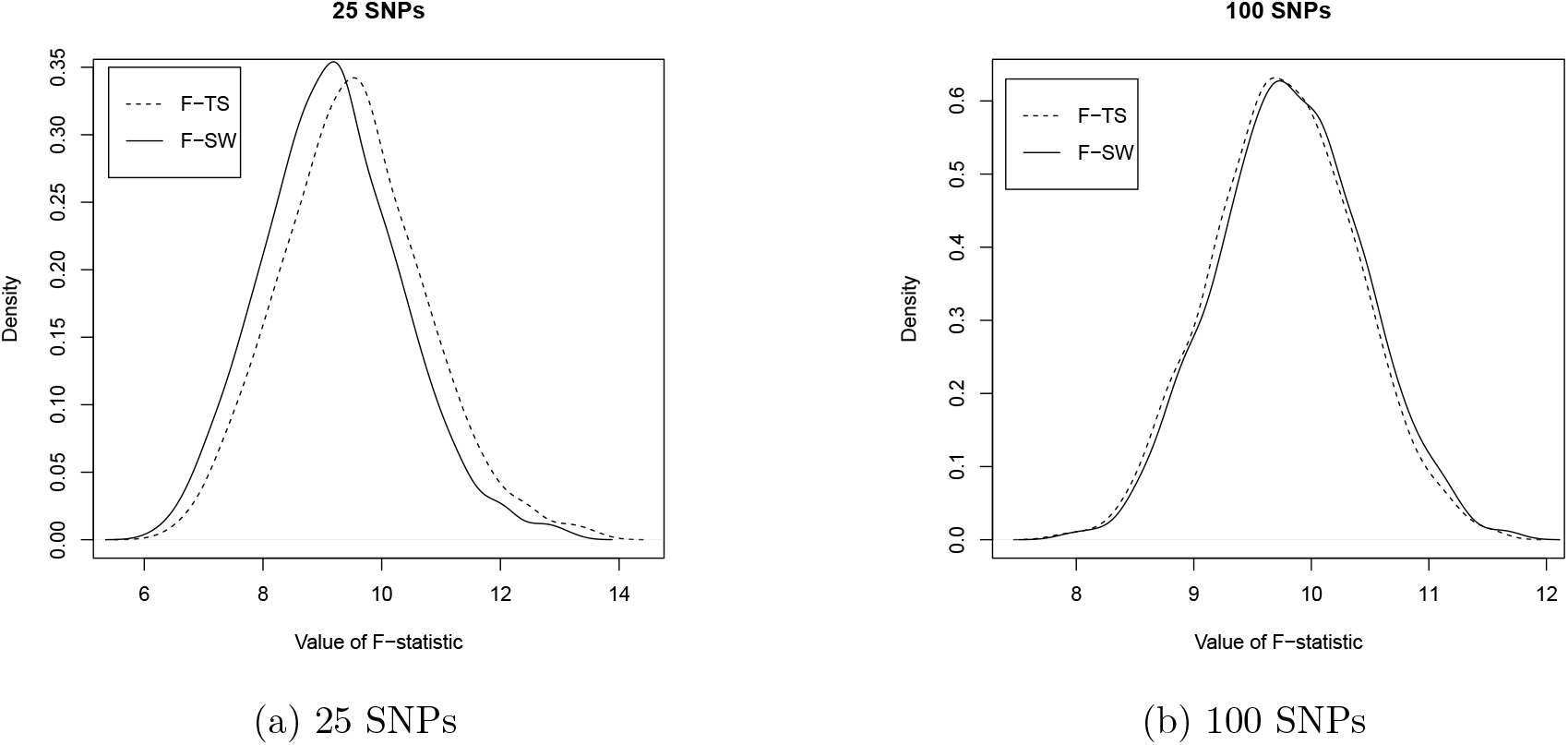
Density of *F_SW_* and *F_TS_*

### Critical values

Comparing this statistic to standard critical values from the F-distribution provides a test for a lack of identification. However, even if the genetic instruments explain some of the variation in the exposure they could still be ‘weak’. In this case the estimates obtained from the MVMR estimation could still be considerably biased. The one sample conditional F-statistic (*F_SW_*) has the same distribution as the Stock-Yogo weak instrument test^6^. Therefore we can apply its weak instrument critical values to identify weak instrument bias for univariable and multivariable two-sample MR^5,6,7^. The weak instrument critical values derived by Stock and Yogo (2005) for the bias of the 2SLS estimator relative to the OLS estimator have a non-central *χ*^2^ distribution, with *L* degrees of freedom and a non-centrality parameter that is a function of *L* and *K*, divided by *K*. These critical values are derived under the definition that the instruments are weak when the bias of the IV estimator relative to the OLS estimator is at least 10%. The measure of relative bias used is the squared bias of the IV estimator (*β_IV_*) relative to the squared bias of the OLS estimator (*β_OLS_*). This is given by the equation;

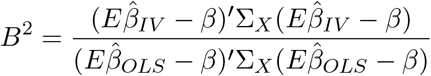

Where 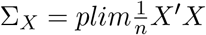 and *X* here represents the *n* × *K* matrix of all of the exposures included in the estimation. Calculating the bias in this way standardises the exposures X so they are orthogonal and have unit standard deviation. However it mean that the bias of the estimated effect of any particular exposure may differ from 10%. If *F_TS_* is larger than the relevant Stock-Yogo critical value we can reject the null hypothesis that the exposure is only weakly predicted by the instruments. These critical values have only been derived for models including up to 30 instruments, therefore in Table 1 we provide critical values for a larger range of instruments to test for a 5%, 10% or 20% relative bias. These critical values are often approximated to a rule of thumb of *F* > 10 to test a null hypothesis that the bias is at least 10% of the bias of the OLS estimator. The critical values given above also show that the rule of thumb of 10 is slightly smaller than the true critical value for this test and would lead to the null hypothesis being rejected more frequently. The two sample *F_TS_* statistic tests the bias of the model as a whole, this means that the sign of the bias of an individual causal parameter may differ from that of the model’s bias, which is averaged across all of its constituent parameters.

**Table 1:** Critical values for conditional weak instrument tests.

| $k_Z$ | Relative bias | | |
| --- | --- | --- | --- |
|  | 5% | 10% | 20% |
| 25 | 21.37 | 11.44 | 6.19 |
| 50 | 21.26 | 11.14 | 5.86 |
| 100 | 21.02 | 10.84 | 5.64 |
| 200 | 20.79 | 10.61 | 5.46 |
| 300 | 20.62 | 10.52 | 5.38 |
| 400 | 20.56 | 10.45 | 5.32 |
| 500 | 20.50 | 10.40 | 5.29 |

## 3 Weak Instrument Robust Two-sample MVMR

### Estimation in the presence of weak instruments

In the presence of weak instruments, standard inverse variance weighted estimation of the MVMR mode, which we refer to as MVMR-IVW, is biased. The reasons for this will be explained in more detail below. The LIML estimator has previously been proposed as an alternative estimator for individual-level MR as it is less biased when there are many weak instruments^15^. In the two-sample summary data setting, Bowden and colleagues ^16^ and Zhao and colleagues^17^ show that weak instruments can be effectively mitigated through minimisation of an appropriate heterogeneity statistic using weights that account for the variance of the SNP-exposure associations is analagous to one-sample LIML estimation. It gives results that are substantially less biased than conventional regression based IVW estimates in the presence of a non-zero causal effect. The weak instrument robust estimation proposed by Bowden and colleagues can be extended to the MVMR setting as a minimisation of;

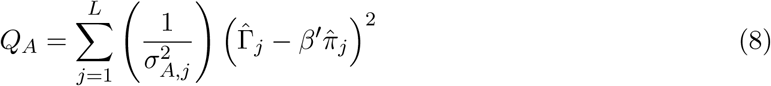

over *β*, Where *β* is a vector of causal parameters (to be estimated), 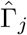 is the estimated effect of SNP *j* on the outcome, 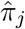 is a vector of effects of SNP *j* on each exposure included in the estimation (i.e. a column of the matrix Π) and;

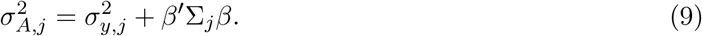

Here, 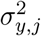 is the variance of the estimated effect of the SNPs on the outcome, and Σ_*V,j*_ is the variance-covariance matrix defined in equation (5). This is equivalent to minimisation of the *Q_A_* statistic to test for heterogeneity described in Sanderson et al 2019^4^ extended to a model with more than two exposures. We label estimates for *β* obtained in this manner as 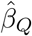. The standard MVMR-IVW estimate is vunerable to weak instrument bias because instead of minimising *Q_A_* in (8) using the full weights defined in (9) it incorrectly assumes that 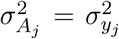. This ignores the component of variation from *β′*Σ_*j*_*β* and is only valid if either all elements of *β* are zero or Σ_*j*_ is negligable by comparison to 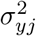.

### Testing for pleiotropy in the presence of weak instruments

Horizontal pleiotropy - where genetic variants influence the outcome through multiple phenotypes can lead to a violation of the IV assumptions if they are not included as exposures in the MVMR estimation. Under the assumption that not all the SNPs included in the estimation have a pleiotropic effect on the outcome through the same pathway, this will lead to greater variation in the estimated causal effect of the exposures on the outcome than would be expected by chance. This excess heterogeneity can be reliably tested for using the minimised *Q_A_* statistic. More formally if all SNPs used in the MVMR analysis are valid instruments, in the sense that they identify a common set of causal parameters *β*, we would expect the *Q_A_* statistic in (6) evaluated at 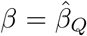 to follow a Chi-squared distribution with L-K degrees of freedom. Crucially, the test is exact in the sense that it will achieve its nominal type I error rate, even in the presence of weak instruments.^18^. The standard Q-statistic used to generate the MVMR IV estimate by setting 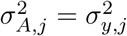, referred to here as *Q_IVW_*, will generally have an inflated type 1 error rate (i.e. will detect pleiotropy too often when none is present) unless all *β′*Σ_*j*_*β* terms are negligible.

### Estimation in the presence pleiotropy and weak instruments

Estimation of *β* through minimisation of (8) will give estimates of the direct effect of each exposure on the outcome that are robust to weak instruments. However, these estimates will still be biased in the presence of pleiotropy. In order to account for heterogeneity due to pleiotropy, we extend the estimation of *β* by adding a pleiotropy variance parameter *τ*^2^ to the multivariable Q estimation and finding the joint value of (*β, τ*^2^) which minimises;

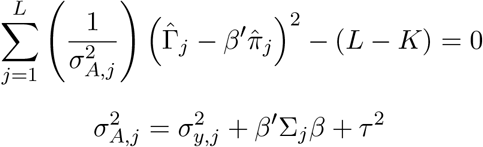

subject to;

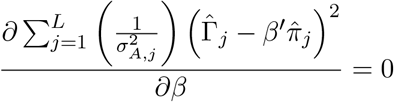

We refer to the causal estimates derived in this way as 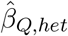 This is a extension of the method described in^16^ for univariable MR to the MVMR setting. Although it will account for balanced pleiotropy, where there are equal positive and negative pleiotropic effects of the SNPs on the outcome, this method of estimation will not adjust for directional pleiotropy where the pleiotropic effects of the SNPs on the outcome all, or mostly, act in one direction to either increase or decrease the outcome. However, it is possible to look at the individual contribution of each SNP to *Q_A_* to identify the largest outliers. If a small number of SNPs are observed to have a large effect on *Q_A_* they can potentially be removed as a sensitivity analysis and the MVMR model re-estimated without them.

### Obtaining confidence intervals for estimated effects

Estimation of *β* and *τ*^2^ through minimisation of *Q_A_*, does not provide readily available and reliable standard errors. We therefore suggest that standard errors are obtained, and confidence intervals calculated, through a Jackknife procedure.

We propose the use of Jackknife rather than a bootstrap as with a moderate number of SNPs the repeated sampling in a bootstrap can lead to very weak instruments in any particular iteration even when the model has relatively strong instruments as a whole. A jackknife procedure estimates the model leaving out each SNP in turn and then calculates the standard deviation of the effect estimate from these results. As each iteration includes all but one of the SNPs and includes each SNP only once this is unlikely to be affected by weak instruments due to the exclusion of some SNPs.

## 4 Estimation of *σ_ij_*

So far we have assumed that the pairwise covariance between a SNPs estimated association with any two exposures is known for all exposures and all SNPs. However, this data is not generally reported by GWAS summary statistics. Similarly it would not be feasible for these studies to report this data due to the large number of potential covariances that could be required for all potential future MVMR analyses. Results from our simulations (given in section 5) show that estimation without these covariances (i.e. imposing the assumption *σ_i,j_* = 0, *i* ≠ *j*) can have large effects on the results obtained, altering the interpretation of the results. Excluding these covariances will give the correct estimation only under the global null (*β* = 0).

Therefore, in this section we suggest three different solutions for dealing with the lack of co-variances in the GWAS summary results in order to estimate *σ_km,j_*: the covariance between 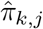 and 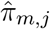 with respect to exposure, *k*, exposure, *m* (*k* ≠ *m*) and SNP *j*.

### Estimate *σ_km,j_* from the individual level data

If some or all of the individual level data that was used in the GWAS to estimate the SNP - exposure associations is available then the covariances for the effect of each SNP on each exposure can be calculated from equation 6.

### Estimate the phenotypic correlation between the exposures from individual level data

The covariance for each SNP can then be approximated as;

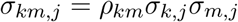

where *ρ_km_* is the correlation between *X_k_* and *X_m_* (or phenotypic correlation). As each SNP explains only a small proportion of the variation in each exposure, the covariance between the exposures provide a very good approximation for the covariance between the error terms in the SNP exposure associations. Although ideally this information would be calculated from the data used for the GWAS study, *ρ_km_* could also be estimated from only part of the data used in the GWAS or from an alternative dataset which is thought to have a similar structure.

### Estimate the effect of the SNPs on each exposure from separate samples

Estimating the effect of the SNPs on each exposure in this manner means that the covariances will be zero and so excluding this information will not affect the statistics calculated. For an MVMR analysis involving *K* exposures, this would require *K* + 1 separate samples and so is likely to only be practicable in a limited number of cases.

In any given scenario some of these solutions may be impossible (due to a lack of data) and of the solutions that are possible, one may be the most reasonable. We suggest that estimation of *ρ_km_* from phenotypic data, from which the appropriate covariances can then be calculated, is likely to be the most feasible and appropriate approach in many cases.

## 5 Simulation Results

To illustrate the methods presented so far give here results from simulating and fitting MVMR models with 200 SNPs and either 2 or 3 exposures.

### MVMR model with two exposures

Firstly, we simulated a MVMR model with 2 exposures and 200 SNPs. The SNP-exposure associations where constructed in two ways; firstly so that each exposure was individually and conditionally weakly predicted by the set of SNPs (i.e. weak instruments) and secondly so that the exposures were strongly individually predicted, but weakly conditionally predicted by the set of SNPs (conditionally weak instruments). Conditionally weak instruments were generated by increasing the total strength of the instruments but introducing correlation between the effect of each SNP on each of the exposures. This reflects a scenario where examination of standard F-statistics for each exposure would not identify weak instruments. The exposures were simulated to both have a direct effect on the outcome and balanced pleiotropy was introduced to the model through a direct effect of the SNPs on the outcome. Pleiotropic effects were generated from a normal distribution with zero mean. A confounder of both exposures and the outcome was also included. The covariance parameter *σ_i,j_*, *i* ≠ *j* was estimated from calculation of the phenotypic correlation between *X*_1_ and *X*_2_ as described in section 4. The set up of this model is illustrated in Fig. 3 and results from the simulation are given in Table 2. Results for the same model without the pleiotropic effect of the SNPs on the outcome are given in Supplementary Table S.1.

**Figure 3:**
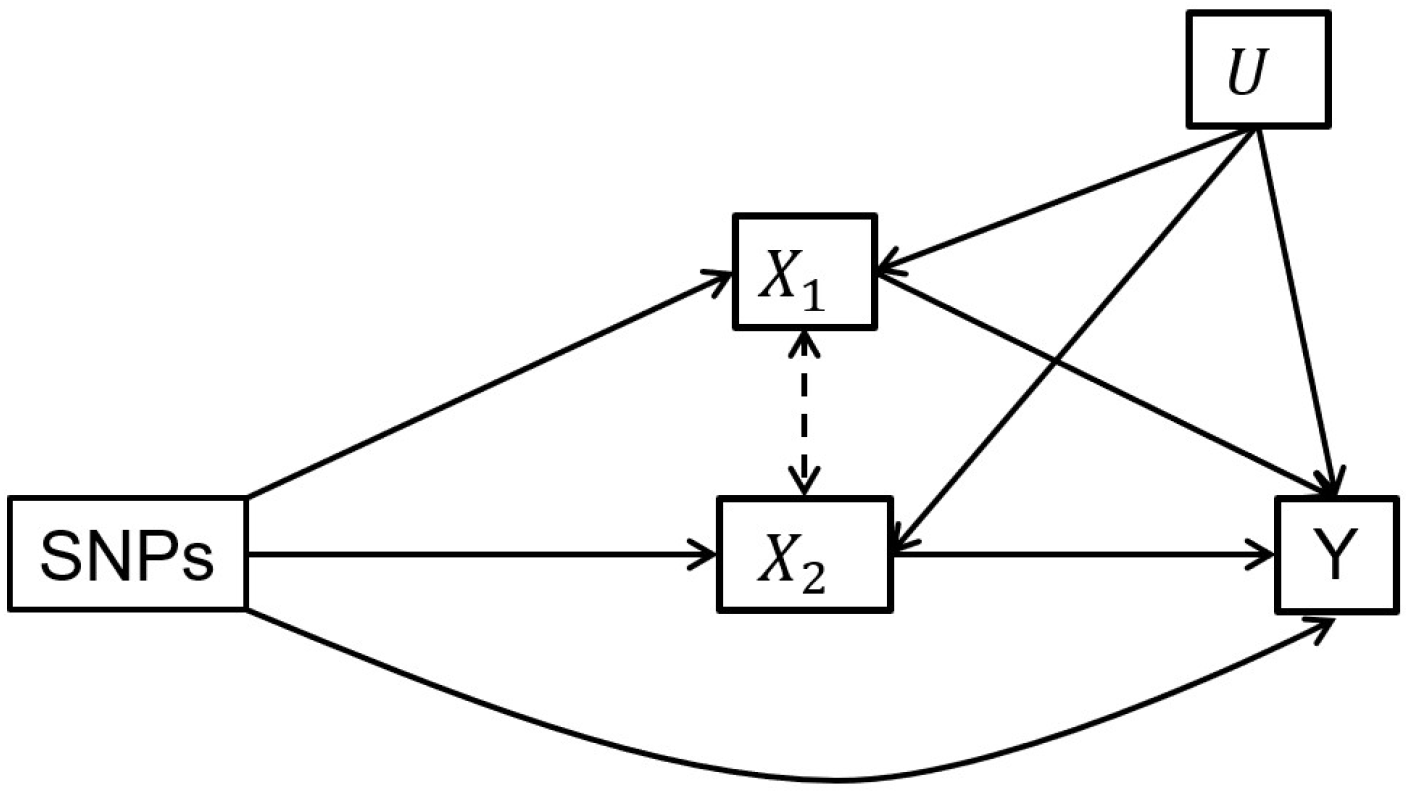
Model simulated in Table 2

**Table 2:**
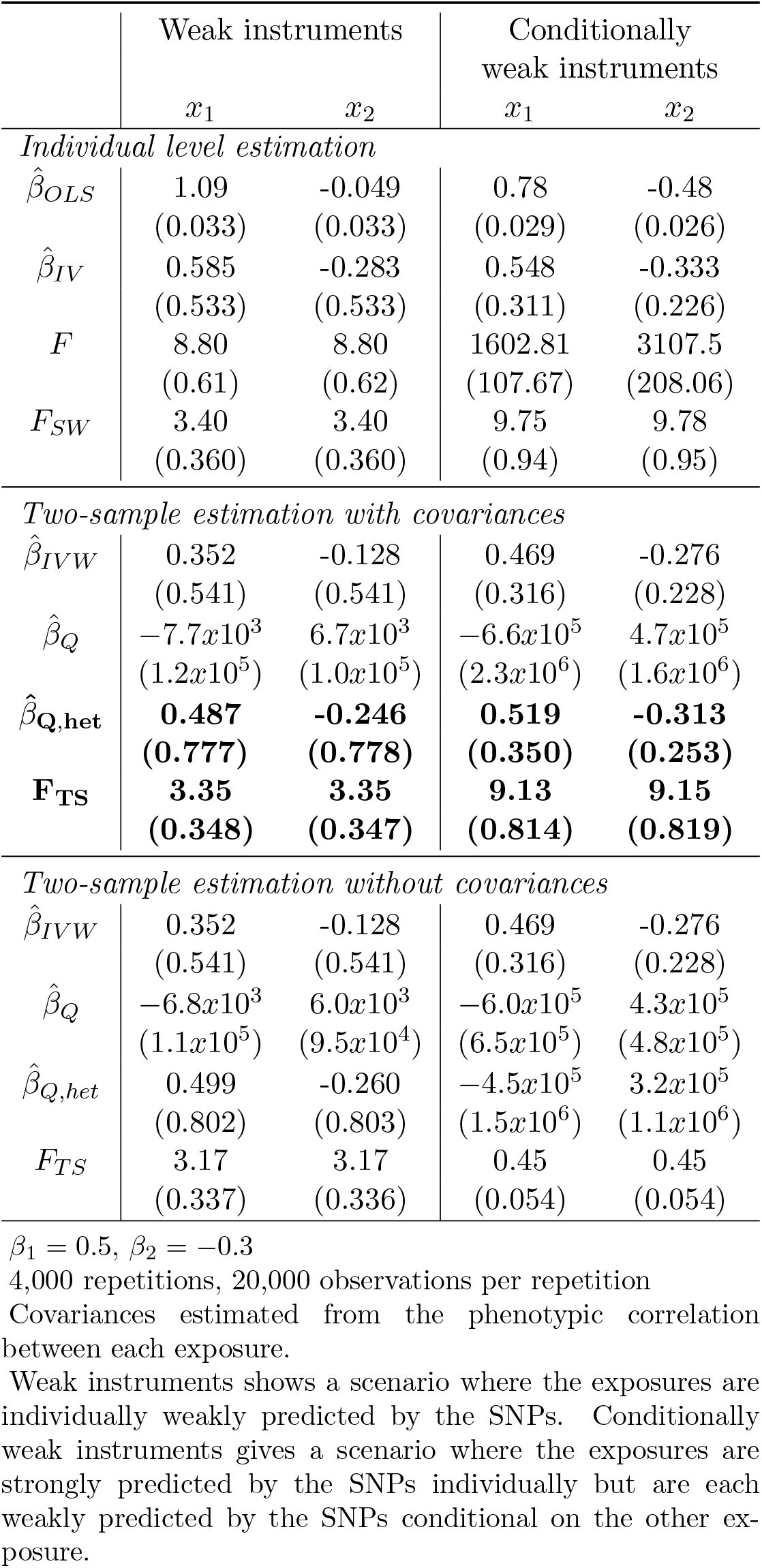
Simulation results for models with heterogeneity: 2 exposures, 200 SNPs

Results from this simulation show that the two-sample conditional F statistic *F_TS_* reliably estimates the strength of the instruments and is equivalent to the conditional F statistic calculated from the individual level data *F_SW_* when the correlation between the exposures is used to estimate the covariance between the effect of each SNP on each exposure. These results also show that although 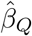 does not reliably estimate the effect of the exposure on the outcome in the presence of balanced of pleiotropy, 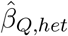 which allows for this additional heterogeneity does. This decrease in bias in 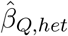 compared to 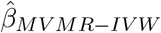 when the instruments are weak comes at the cost of increased standard errors, reflecting the (true) lower level of information in the model. Supplementary Table S.1 shows that allowing for heterogeneity when it is not present does not increase the standard error of the 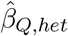 estimates relative to the standard error of the 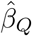 estimate. Table 2 also gives *F_TS_* and *β_Q,het_* estimated without accounting for *σ_km_*, labelled *F*_*TS*,0_ and 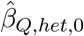 respectively. This imposes the assumption that *σ_km_* = 0 *k* ≠ *m* but not the assumption that 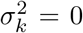 and so is a point between standard MVMR-IVW estimation and 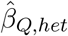. These results also show that in the presence of conditionally weak instruments, when there is correlation between the effect of the SNPs on each exposure, if these correlations are not taken into account *F*_*TS*,0_ does not reliably test the strength of the instruments and 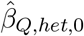 produces biased estimates of the effect of each exposure on the outcome.

### Three exposure model

Next we simulated summary data for three exposures and 200 SNPs. Each of the exposures was simulated to have a direct effect on the outcome. The effect of the SNPs on the first and third exposures were correlated, so that the third exposure was only weakly predicted by the SNPs conditional on the first exposure (and therefore the first exposure is weakly predicted conditional on the third exposure). This set up means that when only the first two exposures are included in the estimation there is directional pleiotropy present, however when all three exposures are included there is potential weak instrument bias. The model under which the data was generated is illustrated in Fig. 4 and results are given in Table 3.

**Figure 4:**
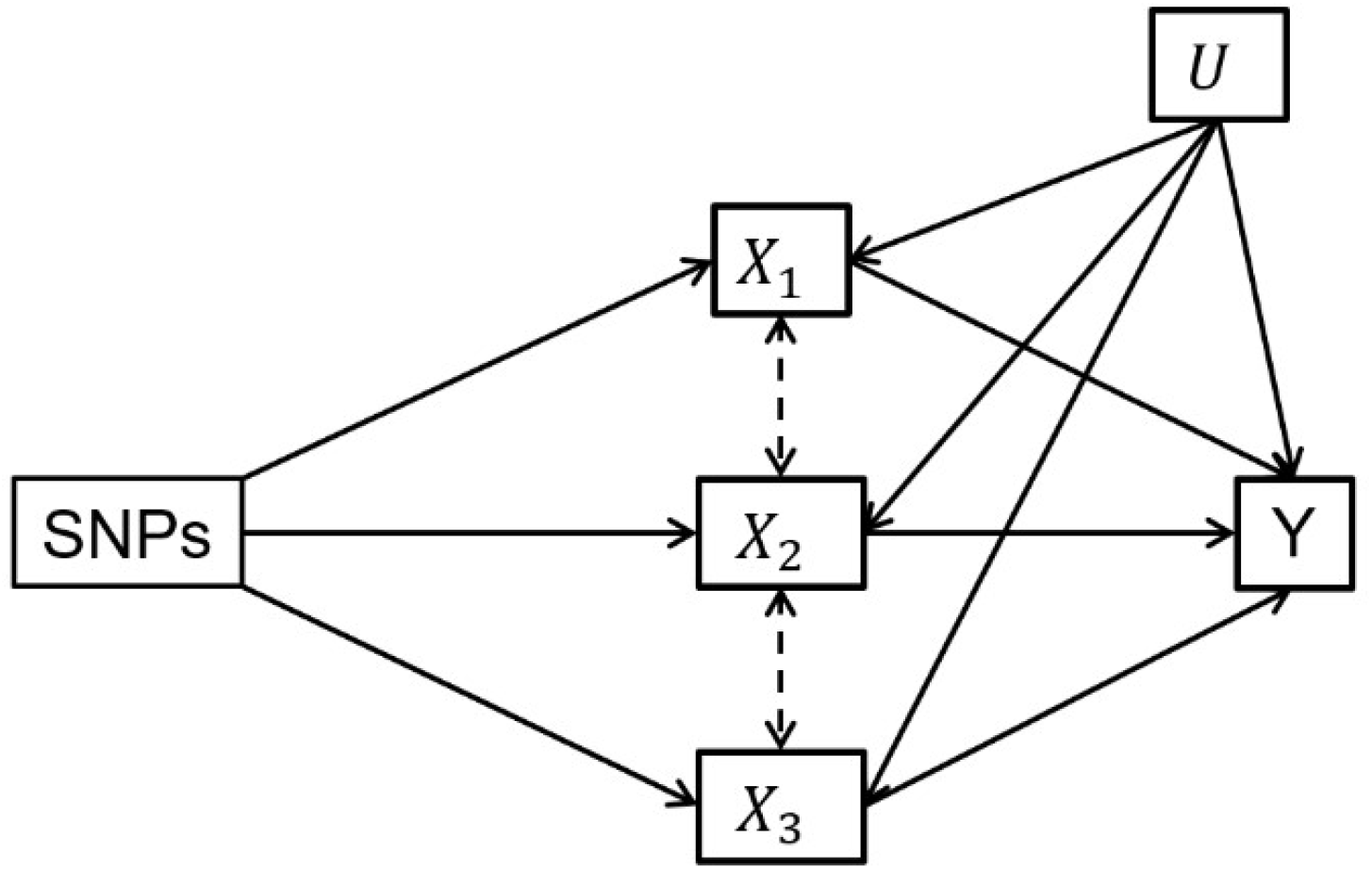
Model simulated in Table 3

**Table 3:** Simulation results for a model with three exposures.

|  | Two exposures<br>included in estimation |  | Three exposures<br>included in estimation |  |  |
| --- | --- | --- | --- | --- | --- |
| | $x_1$ | $x_2$ | $x_1$ | $x_2$ | $x_3$ |
| *Individual level estimation* |  |  |  |  |  |
| $\hat{\beta}_{OLS}$ | 0.837<br>(0.020) | -0.065<br>(0.019) | 0.667<br>(0.020) | -0.176<br>(0.018) | 1.418<br>(0.012) |
| $\hat{\beta}_{IV}$ | 0.626<br>(0.018) | -0.244<br>(0.018) | 0.466<br>(0.017) | -0.314<br>(0.014) | 0.912<br>(0.064) |
| $F$ | 236.2<br>(15.43) | 235.13<br>(13.67) | 236.22<br>(15.43) | 235.13<br>(13.67) | 14.50<br>(0.89) |
| $F_{SW}$ | 78.49<br>(7.10) | 78.37<br>(6.98) | 6.62<br>(1.13) | 19.81<br>(5.38) | 3.17<br>(0.29) |
| *Two-sample estimation with covariances* |  |  |  |  |  |
| $\hat{\beta}_{IVW}$ | 0.611<br>(0.021) | -0.228<br>(0.021) | 0.523<br>(0.027) | -0.277<br>(0.021) | 0.502<br>(0.101) |
| $\hat{\beta}_Q$ | 0.626<br>(0.022) | -0.246<br>(0.021) | 0.500<br>(0.034) | -0.301<br>(0.024) | 0.703<br>(0.149) |
| $\hat{\beta}_{Q,het}$ | **0.624**<br>**(0.022)** | **-0.246**<br>**(0.021)** | **0.499**<br>**(0.035)** | **-0.301**<br>**(0.024)** | **0.705**<br>**(0.154)** |
| $F_{TS}$ | **45.01**<br>**(2.39)** | **44.97**<br>**(2.35)** | **6.58**<br>**(1.09)** | **17.94**<br>**(4.32)** | **3.23**<br>**(0.29)** |

$\beta_1 = 0.5, \beta_2 = -0.3, \beta_3 = 0.7$
4,000 repetitions, 20,000 observations per repetition
Covariances estimated from the phenotypic correlation between each exposure.

We give results from estimation of the model firstly including only two exposures, *x*_1_ and *x*_2_, and including all three exposures. These results show that when only two exposures are included in the model all methods of estimating *β*_1_ and *β*_2_ are biased by the directional pleiotropy present in the model. When all three exposures are included in the model the MVMR-IVW estimates are biased due to the presence of weak instruments. However, estimation of 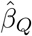 through minimisation of *Q_A_* gives unbiased estimates of the effect of each exposure.

### Heterogeneity Testing

Table 4 gives the rejection rates when using *Q_IVW_* and *Q_A_* to test for pleiotropy for the model considered in Figure 2. In addition, we show rejections rates using a third heterogeneiyt statistic that attemps to improve 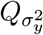 by extending the weights so they take the form 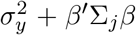. We call this heterogeneity statistic *Q_IVW,up_*. These results show that when there is no heterogeneity the null hypothesis is over rejected by both *Q_IVW_* and *Q_IVW,up_*. Estimation of *Q_A_* accounting using weak instrument robust estimates of 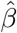 and accounting for the variation in the SNP-exposure association controls the type 1 error and when the null hypothesis is true, i.e. when there is no heterogeneity this test statistic rejects approximately 5% of the time.

**Table 4:** Estimation of *Q_A_*.

|  | Weak instruments |  |  |  | Conditionally<br>weak instruments |  |  |  |
| --- | --- | --- | --- | --- | --- | --- | --- | --- |
| | $\tau^2 = 0$ | | $\tau^2 = 0.5$ | | $\tau^2 = 0$ | | $\tau^2 = 0.5$ | |
|  | Estimate | Rej. Rate | Estimate | Rej. Rate | Estimate | Rej. Rate | Estimate | Rej. Rate |
| $Q_{IVW}$ | 228.37<br>(23.04) | 41.9% | 13366.76<br>(678.27) | 100% | 249.71<br>(24.85) | 75.4% | 13032.82<br>(770.59) | 100% |
| $Q_{IVW,up}$ | 206.20<br>(21.39) | 12.9% | 11645.15<br>(1699.61) | 100% | 201.36<br>(20.53) | 7.6% | 11593.22<br>(1338.27) | 100% |
| $Q_A$ | 197.16<br>(19.83) | 4.5% | 576.93<br>(53.92) | 100% | 197.74<br>(19.85) | 5.2% | 1788.42<br>(224.66) | 100% |

4,000 repetitions, 20,000 observations per repetition
Covariances estimated from the phenotypic correlation between each exposure.

## 6 Application

In this section we illustrate the use of the methods described above through an application to the estimation of the effect of multiple metabolites to age-related macular degeneration (AMD). AMD is disease that causes loss of central vision and is a leading cause of blindness^19^. Elevated lipid serum levels have previously been associated with increased risk of AMD^20^. We use data from a Genome-wide association study (GWAS) of 118 metabolites by Kettunen et al 2016^14^ as our exposure and from a GWAS of AMD as our outcome ^21^. Previous studies have implicated HDL as being causal for AMD^22,23,24^. Our analysis shows that other lipid fractions do not have a causal effect on AMD.

The GWAS data included 118 potential metabolite exposures. For the purposes of illustration we restricted the analysis to 14 metabolites moderately well predicted by a large number of SNPs. From the 150 SNPs included in the data we selected a set of 78 which were associated with at least one of our exposures with an F statistic greater than 5. Table 5 gives MVMR-IVW estimates of all of these metabolites against AMD. As well as the MVMR-IVW estimation results, Table 5 also includes; the mean F-statistic for the SNPs associated with each metabolite (*F_individual_*), the mean F-statistic across all of the SNPs included in the analysis for each metabolite (*F_all_*) and the conditional F-statistic for each metabolite (*F_TS_*). The correlation between the metabolites was not available from the GWAS data used here. We therefore calculated these using external data on the same metabolites from the Avon Longitudinal Study of Mothers and Children (ALSPAC) ^25,26^. A description of the ALSPAC study is given in the supplementary material. The F-statistics and conditional F-statistics presented show that although each metabolite is strongly predicted by the SNPs associated with it, and most have a *F_all_* > 10, the conditional F-statistics for each exposure are very small and therefore the effect estimates are subject to weak instrument bias.

**Table 5:**
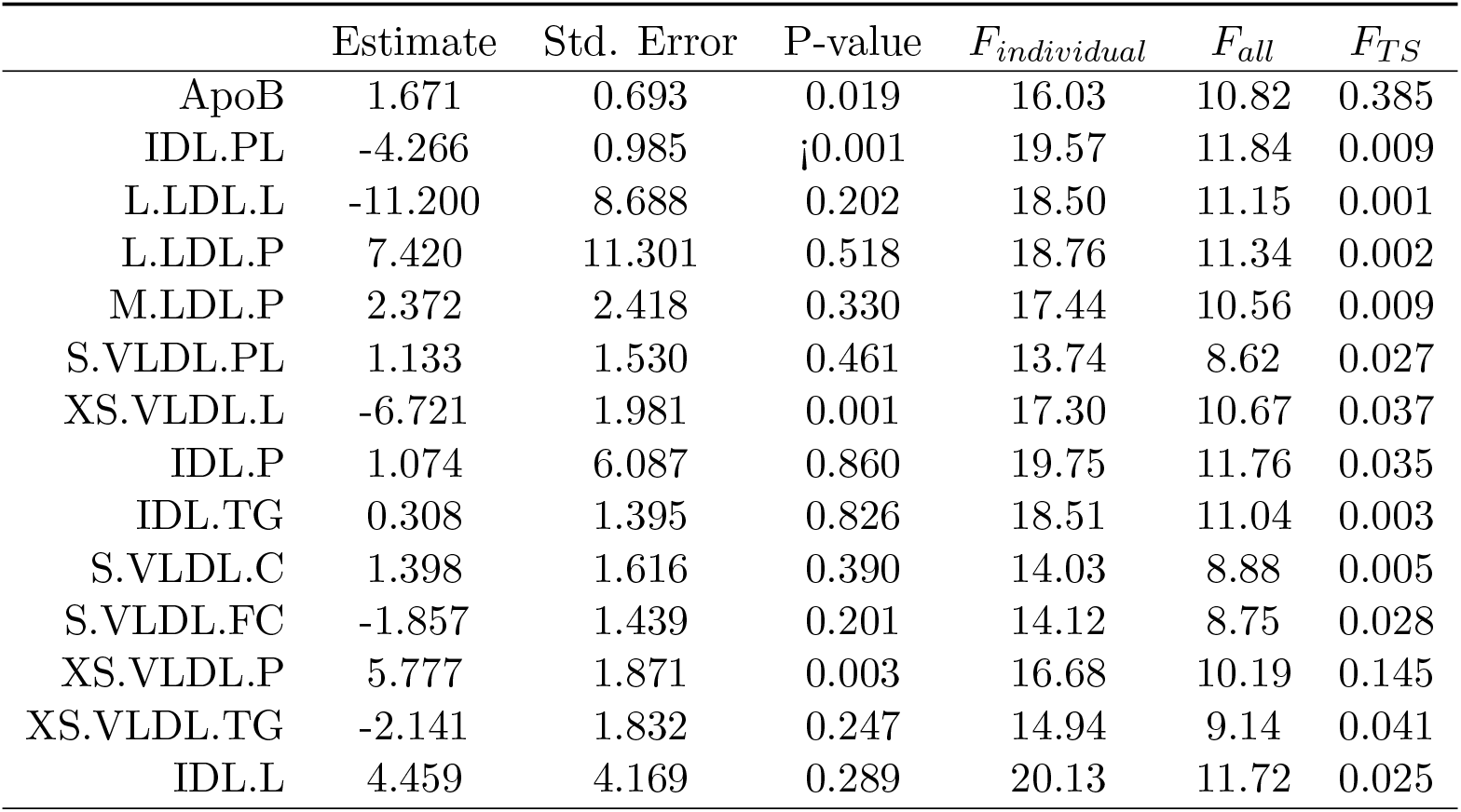
MVMR estimates of a range of metabolites on AMD

This set of metabolites can be divided into four groups, IDL, LDL, small VLDL and very small VLDL. In Table 6 we present the results for the calculation of the F-statistics *F_all_* and *F_TS_* for each of these groups. They show that, with the exception of IDL.PL and small VLDL, the SNPs are too conditionally weak to predict all exposures each of these groups. Further investigation (not shown) indicated that the SNPs are too conditionally weak to predict any pair of SNPs from any group other than small VLDL or IDL.PL.

**Table 6:** F and conditional F statistics for groups of traits

|  | IDL |  | LDL |  | Small VLDL |  | Very Small VLDL |
| --- | --- | --- | --- | --- | --- | --- | --- |
| | $F$ | $F_{TS}$ | $F$ | $F_{TS}$ | $F$ | $F_{TS}$ | |
| ApoB |  |  |  |  |  |  |  |
| IDL.PL | 16.05 | 8.744 |  |  |  |  |  |
| L.LDL.L |  |  | 17.58 | 0.019 |  |  |  |
| L.LDL.P |  |  | 17.83 | 0.023 |  |  |  |
| M.LDL.P |  |  | 16.55 | 0.063 |  |  |  |
| S.VLDL.PL |  |  |  |  | 12.38 | 11.65 |  |
| XS.VLDL.L |  |  |  |  |  |  | 14.64 0.0174 |
| IDL.P | 16.12 | 0.010 |  |  |  |  |  |
| IDL.TG | 14.97 | 0.012 |  |  |  |  |  |
| S.VLDL.C |  |  |  |  | 12.42 | 4.75 |  |
| S.VLDL.FC |  |  |  |  | 12.51 | 5.39 |  |
| XS.VLDL.P |  |  |  |  |  |  | 12.50 0.196 |
| XS.VLDL.TG |  |  |  |  |  |  | 13.99 0.176 |
| IDL.L | 16.08 | 0.009 |  |  |  |  |  |
| SNPs | 54 |  | 46 |  | 50 |  | 53 |

We therefore considered including one metabolite from each group as our exposures. For this set of exposures we calculated the conditional F-statistics, *F_TS_* and the MVMR-IVW estimates. These results are given in Table 7. Although the exposures here are jointly moderately strongly predicted by the set of SNPs the conditional F-statistics for each exposures are still between 4.2 and 8.3 indicating that there may be some weak instrument bias. In Table 8 we re-estimate our MVMR model using the weak instrument robust methodology presented earlier. These results show that the initial MVMR including all of the metabolites could give misleading results due to weak instrument bias. *Q_A_* for this model is 118, the critical value at a 5% level of significance for a chi-squared distribution with 64 degrees of freedom is 84.7. It therefore indicates potential pleiotropy and we consider the 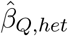 to be the most appropriate estimates in this case. The results from this analysis suggest that none of the metabolites considered are causally associated with AMD but that the standard MVMR IVW estimates for the final model were biased due to both weak instruments and pleiotropic effects of the SNPs on the outcome. This null result is consistent with other results using an alternative method to analyse the same data which found that HDL (not included in this analysis) was the only lipid fraction type that was causally associated with AMD^24^.

**Table 7:** MVMR-IVW estimates of a range of metabolites on AMD including only strongly predicted traits

| | Estimate | Std. Error | P-value | $F_{all}$ | $F_{TS}$ |
| --- | --- | --- | --- | --- | --- |
| XS.VLDL.P | -0.778 | 0.958 | 0.420 | 11.26 | 4.23 |
| S.VLDL.PL | 0.051 | 0.347 | 0.385 | 9.48 | 5.68 |
| L.LDL.L | 0.356 | 0.231 | 0.154 | 12.19 | 8.22 |
| IDL.TG | 0.067 | 0.761 | 0.969 | 12.21 | 6.15 |

69 SNPs

**Table 8:** MVMR-IVW estimates of a range of metabolites on AMD including only strongly predicted traits

| | Updated MVMR-IVW | | | $\hat{\beta}_Q$ | | | $\hat{\beta}_{Q,het}$ | | |
| --- | --- | --- | --- | --- | --- | --- | --- | --- | --- |
|  | Est. | Std. Error | p-value | Est. | Std. Error | p-value | Est. | Std. Error | p-value |
| XS.VLDL.P | -0.267 | 1.051 | 0.799 | -5.008 | 3.774 | 0.0185 | -2.071 | 1.447 | 0.152 |
| S.VLDL.PL | -0.094 | 0.346 | 0.786 | 0.957 | 0.940 | 0.309 | 0.300 | 0.528 | 0.570 |
| L.LDL.L | 0.328 | 0.225 | 0.145 | 1.534 | 0.645 | 0.017 | 0.728 | 0.613 | 0.235 |
| IDL.TG | -0.311 | 0.812 | 0.702 | 2.490 | 2.614 | 0.341 | 0.803 | 1.437 | 0.576 |

69 SNPs

## 7 Software

We have written an R package MVMR which facilitates the implementation of MVMR estimation and corresponding sensitivity analyses. The package requires summary data on instrument-exposure and instrument-outcome associations, as well as information on the pairwise covariances of the error in the estimated association between each SNP and each pair of exposures. As these covariances are often not available the software can be implemented in three ways; estimating the covariances from individual level data, approximating the covariances from the phenotypic correlation between the exposures or assuming that these covariances are zero.

### Workflow

Fitting and interpreting MVMR using the methods described in this paper, including tests for instrument strength and horizontal pleiotropy, is performed using a five-step procedure. Initially, summary data should be provided, including a covariance matrix for the effect of the genetic variants on each exposure. As such covariances are not conventionally reported in publicly available data, two functions `snpcov_mvmr()` and `phenocov_mvmr()` can be used to generate the covariance matrix. The `function snpcov_mvmr()` estimates the covariance terms directly from individual level data, whilst `phenocov_mvmr()` uses the phenotypic correlation and summary data (input by the user) to generate estimates of the covariances.

As a second stage, the summary data is reformatted using the function `format_mvmr()` into a data frame which is subsequently used as the input for estimation and sensitivity analyses. We then provide the functions `strength_mvmr()` to evaluate instrument strength using the two sample conditional F-statistic described in Section 2. Tests for horizontal pleiotropy are performed using `pleiotropy_mvmr()`, performing both standard and Q-minimisation approaches simultaneously (see section 3 for more details). Finally, causal effects can be estimated using two different approaches; fitting an inverse variance weighted (IVW) MVMR model using `ivw_mvmr()`and minimising the Q-statistic allowing for heterogeneity using `qhet_mvmr()`. Each step in the MVMR workflow is illustrated in Figure 5. The MVMR package is available to download at https://github.com/WSpiller/MVMR/. The package also includes a detailed tutorial demonstrating functionality of the package in an analyses of the effects of lipid fractions upon systolic blood pressure using data from the Global Lipids Genetics Consortium and UK Biobank.

**Figure 5:**
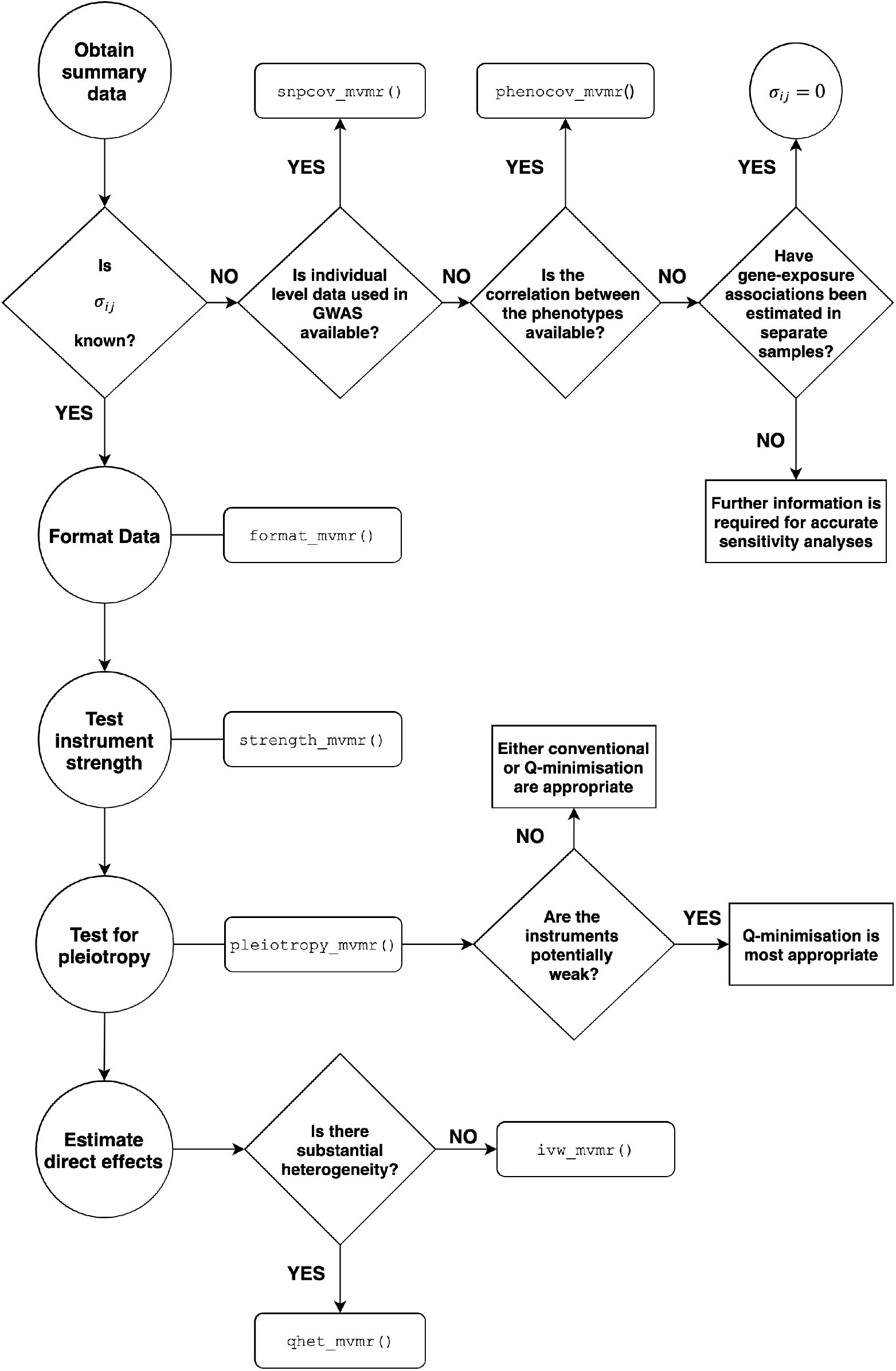
Workflow for MVMR R package

## 8 Discussion

In this paper we develop a general statistical framework for conducting two sample MVMR analyses for an arbitrary number of exposures in the presence of weak instrument bias and pleiotropy. The methods presented here give ways to test for weak instruments in two-sample MVMR and to robustly test for heterogeneity due to pleiotropy in the presence of moderately weak instruments. We additionally give a method to estimate causal effects in the presence of moderately weak instruments which is robust to balanced pleiotropy.

Weak instruments are a potential issue in many applications where estimating direct effects of multiple exposures using MVMR is preferred over univariable MR analyses, which are thought to be likely to be affected by directional pleiotropy^27,28,29,30,31^. MVMR approaches are also used to gague the extent to which one exposure mediates the effect of another on the outcome ^32,33^.Any application of MVMR will be biased by conditionally weak instruments and, as illustrated by our application, this can occur even when the genetic variants strongly predict each exposure individually. Therefore, the methods presented here are important as they provide a way to identify and correct for weak instruments in two-sample MVMR estimation.

The *F_TS_* statistic described here is calculated using estimates of 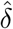 calculated from an IVW estimation of the effect of 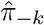 on 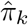. An alternative method of estimation, equivalent to that described for estimation of *β*, is to directly minimise its constituent 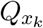 to obtain LIML estimates for *δ* ^16,17^. Whilst this procedure enacted on the *Q_A_* statistic furnishes attractive, weak instrument robust causal estimates, initial simulation results (not reported here) showed limited benefit of estimating *δ* in this way therefore we did not investigate potential implementation further.

There are a number of limitations to this work. The test statistic and weak instrument robust estimation requires an estimate of the covariance between the error in the estimated effect of each SNP on each exposure. Although this data is generally not available we propose a method to estimate it, using the phenotypic correlation between the exposures, which can be used to obtain a reasonable approximation if the relevant covariance when each SNP only explains a small proportion of each exposure. We hope that our work will influence future GWAS consortia to release this information as a matter of course, in order to enable the straightforward application of MVMR methods going forward.

Another weakness of the test statistics provided here is the lack of standard errors for the point estimates of the direct effect of each exposure. We propose using a jackknife to estimate these standard errors. This does however make the estimation of these statistic more computationally intensive than would the case if the standard errors could be calculated analytically.

The weak instrument robust point estimates are robust to weak instruments but cannot produce reliable estimates when instruments become very weak or if only a small number of SNPs are available. Although we show this method works with moderately weak instruments it is not clear exactly how weak is too weak, or indeed how few instruments are too few, to produce either reliable point estimates or heterogeneity statistics. Gaining a more precise understanding of these questions is a topic for further research.

Although we propose weak instrument robust estimation, if the weak instruments are limited to only a small number of the exposures in the model an alternative approach may be to drop exposures (one at a time) until the conditional F-statistics show that all of the exposures are strongly predicted by the SNPs. This would however need to be considered carefully as if dropping an exposure has the potential to introduce directional pleiotropy into the estimation biasing the resulting effect estimates. The choice of approach to take would depend on the number of SNPs and exposures in the estimation and the relationship between the exposures as well as how weak the SNPs are as instruments. As illustrated by our application these approaches could be combined, excluding exposures until instrument strength is high enough to reasonably apply the weak instrument robust methods. The choice approach needs to be considered on a case by case basis.

Additionally although our final estimation *β_Q,het_* is robust to balanced pleiotropy it will still give biased estimates in the presence of unbalanced or directional pleiotropy. Multivariable MR Egger^13^, has been proposed as a method for obtaining reliable MVMR estimates in the presence of directional pleiotropy. Extending this approach to account for weak instrument bias is another topic of further research.

#### Box 1: Summary of statistics discussed in this paper

**Instrument strength statistics;**

*F* - Measure of the strength of the instruments to predict one exposure. Applies to individual or summary level data and to univariable or multivariable MR estimation.

*Conditional F-statistic F_SW_* - Measure of the strength of instruments to predict one exposure conditional on the other exposures included in the estimation. Applies to multivariable MR estimation with individual level data.

*Conditional F-statistic F_TS_* - Measure of the strength of instruments to predict one exposure conditional on the other exposures included in the estimation. Applies to multivariable MR estimation with summary data.

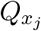 - A *Q*-statistic from which *F_TS_* is calculated.

**Heterogeneity statistics;**

*Q_IVW_* - A heterogeneity test for MVMR that uses the IVW point estimates and does not account for the uncertainty in the estimated SNP-exposure associations. This test over rejects the null in the presence of weak instruments.

*Q_IVW,up_* - A heterogeneity test for MVMR that uses the IVW point estimates but accounts for the uncertainty in the estimated SNP-exposure associations. This test over rejects the null in the presence of weak instruments, but to a lesser extent that *Q_IVW_*.

*Q_A_* - A heterogeneity test for MVMR that is robust to weak instruments, in the sense that it has the appropriate type 1 error rate in the presence of weak instruments.

**Estimation statistics;**

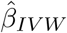 - Estimates of the causal effect of each exposure on the outcome, estimated using standard inverse variance weighting.

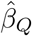 - Estimates of the causal effect of each exposure on the outcome, estimated through minimisation of *Q_A_*. Robust to weak instruments.

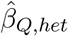 - Estimates of the causal effect of each exposure on the outcome, estimated through minimisation of *Q_A_* with an additional parameter to account for heterogeneity. Robust to weak instruments and pleiotropy.

#### Box 2: Recommended tests in Two-sample MVMR

In all two-sample summary data MVMR estimation two statistics should be calculated;

1. Conditional F statistics, *F_TS_*, for each exposure. These test the strength of the genetic variants to predict each exposure in the multivariable mode. *F*_*TS*_ < 10 suggests potential weak instrument bias in the MVMR estimation.
2. A Q-statistic for heterogeneity, *Q_A_*, for the model. Rejection of *Q_A_* using standard significant levels (e.g. *p* < 0.05) indicates potential pleiotropy in the form of excessive heterogeneity in the MVMR model. However, this test will often reject in the presence of weak instruments.

If weak instruments are detected, i.e. any of the *F*_*TS*_ values are less than 10, IVW MVMR estimates are potentially biased. When large numbers of SNPs are available this can be corrected through;

3. Estimating 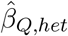 for each exposure This method gives estimates of the direct effect of each exposure on the outcome that are robust to (moderately) weak instruments.
4. An updated *Q_A,min_* which minimises the *Q* statistic over *β_Q_*. This test provides a test for heterogeneity that has the correct size in the presence of weak instruments. Rejection of *Q_A,min_* using standard significant levels (e.g. *p* < 0.05) indicates potential pleiotropy in the MVMR model even in the presence of moderately weak instruments.

All of these tests and estimation statistics are provided in the MVMR R package.

## Code availability

The code used to conduct the simulations and applied analysis is available at https://github.com/eleanorsanderson/MVMRweakinstruments. The MVMR package is available at https://github.com/WSpiller/MVMR/.

## Author contributions

ES and JB devised the project. ES conducted the analysis and wrote the first draft of the paper. WS developed the software package. All authors reviewed and approved the final version.

## Acknowledgements

We are grateful for helpful discussions with Frank Windmeijer and Zoltan Kutalik during the development of this paper.

We are extremely grateful to all the families who took part in the ALSPAC study, the midwives for their help in recruiting them, and the whole ALSPAC team, which includes interviewers, computer and laboratory technicians, clerical workers, research scientists, volunteers, managers, receptionists and nurses.

