## Supplementary material for "Testing and Correcting for Weak and Pleiotropic Instruments in Two-Sample Multivariable Mendelian Randomisation"

### Testing for Weak Instruments in Two Sample Summary Data Multivariable Mendelian Randomisation
### **Supplementary Material**

Eleanor Sanderson[1,2], Wes Spiller[1,2] and Jack Bowden[1,3]

March 2020

#### 1 Equivalence of $F_{TS}$ and $F_{SW}$

The individual level conditional F-statistic derived by Sanderson and Windmeijer (2016), $F_{SW}$, has been shown elsewhere to be an F-test version of the Basmann statistic for overidentification [1,2,3,4]. Here we show that $F_{TS}$ is the two-sample equivalent of the same Basmann statistic and therefore it follows that $F_{TS}$ is equivalent to $F_{SW}$, the equivalent test for individual level data. This equivalence implies that the Stock-Yogo critical values for weak instruments used to test for weak instruments in individual level data are also the appropriate values to compare $F_{TS}$ to to identify weak instruments in two sample Mendelian randomisation. [5,6,1] Although the effect of any particular SNP on different exposures in the model may be correlated we assume throughout that the SNPs themselves are independent of each other. This is not an unreasonable assumption in the context of two-sample MVMR where results from Genome-wide Association Studies are used for each sample. In this context the SNPs used for the analysis are pruned to remove correlated SNPs as standard procedure ensuring that the instruments used in the analysis are independent.

The estimation considered, in the one sample setting, is;

$$y = X\beta + u$$
$$X = Z\Pi + V$$

Where X is a matrix of $K$ exposures and $Z$ is a matrix of $L$ instruments, $K \leq L$. $\beta$ is a vector of effects of $X$ on y, $\Pi$ is a vector of effects of $Z$ on $X$ and $u$ and $V$ are error terms. We can divide the exposures into one of interest and the others by partitioning $X = [X_1\ X_{-1}]$, $V = [v_1\ V_{-1}]$, $\Pi = [\pi_1\ \Pi_{-1}]$. The first stage individual estimation can then be written as;

$$X_1 = Z\pi_1 + v_1$$
$$X_{-1} = Z\Pi_{-1} + V_{-1}$$

As described in the main text the model considered to test the conditional strength of the instruments to predict an exposure is an IV estimation with the exposure of interest as the dependent variable and all other exposures included as exposures predicted by the instruments. In the one sample setting this can be written as;

$$\begin{aligned} X_1 &= \delta_1 X_{-1} + \epsilon_1 \\ X_m &= \sum_{j=1}^{L} \pi_{mj} G_j + \epsilon_m, \qquad m = 2, ..., K \end{aligned}$$

where $X_1$ is the exposure of interest, $X_{-1}$ is a set of all other exposures in the model, $\delta$ is a vector of effects of $X_{-1}$ on $X_1$ and $\epsilon_1$ and $\epsilon_m$ are random error terms. The Basmann statistic to test for overidentification in this model is then given by;

$$\begin{aligned} B &= \frac{\hat{\epsilon} P_Z \hat{\epsilon}}{\left(\hat{\zeta}' \hat{\zeta}\right)/n} \\ &= \frac{(x_1 - X_{-1}\hat{\delta}) P_Z (X_1 - X_{-1}\hat{\delta})}{\left(\hat{\zeta}' \hat{\zeta}\right)/n} \end{aligned}$$

As $\hat{\epsilon} = X_1 - X_{-1}\hat{\delta}$, where $\hat{\delta}$ is a robust IV estimator of $\delta$ and $\zeta$ is the adjusted error term; $\zeta = \hat{v}_1 - \hat{V}_{-1}\hat{\delta}$. Using; $P_Z = Z(Z'Z)^{-1}Z' = Z(Z'Z)^{-1}Z'Z(Z'Z)^{-1}Z'$, $\hat{\pi}_1 = (Z'Z)^{-1}Z'X_1$ and $\hat{\Pi}_{-1} = (Z'Z)^{-1}Z'X_{-1}$,

$$\begin{aligned} (Z'Z)^{-1}Z'(X_1 - X_{-1}\hat{\delta}) &= (Z'Z)^{-1}Z'X_1 - (Z'Z)^{-1}Z'X_{-1}\hat{\delta} \\ &= \hat{\pi}_1 - \hat{\Pi}_{-1}\hat{\delta} \end{aligned}$$

the Basmann statistic can therefore be written as;

$$\begin{aligned} B &= \frac{(X_1 - X_{-1}\hat{\delta})' Z(Z'Z)^{-1}Z'Z(Z'Z)^{-1}Z'(X_1 - X_{-1}\hat{\delta})}{\left(\hat{\zeta}' \hat{\zeta}\right)/n} \\ &= \frac{(\hat{\pi}_1 - \hat{\Pi}_{-1}\hat{\delta})(Z'Z)(\hat{\pi}_1 - \hat{\Pi}_{-1}\hat{\delta})}{\left(\hat{\zeta}' \hat{\zeta}\right)/n}. \end{aligned}$$

As the instruments are independent by construction $Z'Z$ is a diagonal matrix with $Z_j'Z_j$ as the j'th diagonal element and 0's on the off diagonals the numerator of this expression can be written as;

$$(\hat{\pi}_1 - \hat{\Pi}_{-1}\hat{\delta})(Z'Z)(\hat{\pi}_1 - \hat{\Pi}_{-1}\hat{\delta}) = \sum_{j=1}^{L} Z_j'Z_j \left(\hat{\pi}_{1,j} - \hat{\Pi}_{-1,j}\hat{\delta}\right)^2.$$

Now looking at the denominator; as $\hat{\zeta} = (\hat{v}_1 - \hat{V}_{-1}\hat{\delta})$ this can be written as;

$$\hat{\zeta}'\hat{\zeta} = \hat{v}_1'\hat{v}_1 - 2\hat{\delta}\hat{V}_{-1}'\hat{v}_1 + \hat{\delta}'\hat{V}_{-1}'\hat{V}_{-1}\hat{\delta}$$

Using the definitions given in equation 3 we can write;

$$
\begin{aligned}
\hat{v}_1'\hat{v}_1 &= nZ'\Sigma_1 Z \\
\hat{v}_1'\hat{V}_{-1} &= nZ'\Sigma_{12} Z \\
\hat{V}_{-1}'\hat{V}_{-1} &= nZ'\Sigma_2 Z
\end{aligned}
$$

Where $\Sigma_1^2$, $\Sigma_{12}$ and $\Sigma_2$ are variance-covariance matrices for the error terms in the estimate effect of the SNPs on the exposures. As the SNPs are independent from each other these matrices are all diagonal or block diagonal matrices with zeros on the off-diagonals. The expressions above can therefore be written as;

$$
\begin{aligned}
\hat{v}_1'\hat{v}_1 &= nZ'\Sigma_1 Z = n\sum_{j=1}^{L} Z_j' Z_j \sigma_{1,j}^2 \\
\hat{v}_1'\hat{V}_{-1} &= nZ'\Sigma_{12} Z = n\sum_{j=1}^{L} Z_j' Z_j \Sigma_{12,j} \\
\hat{V}_{-1}'\hat{V}_{-1} &= nZ'\sigma_2^2 Z = n\sum_{j=1}^{L} Z_j' Z_j \Sigma_{2,j}
\end{aligned}
$$

Where $Z_j$ is the $j'th$ element of $Z$, $\sigma_{1,j}$ is the $j'th$ diagonal element of $\Sigma_1$, $\Sigma_{12,j}$ is the $j'th$ block diagonal element of $\Sigma_{12}$ and $\Sigma_{2,j}$ is the $j'th$ block diagonal element of $\Sigma_2$. Substituting these results back into $\hat{\zeta}'\hat{\zeta}$ gives;

$$
\begin{aligned}
\hat{\zeta}'\hat{\zeta} &= \hat{v}_1'\hat{v}_1 - 2\hat{\delta}\hat{V}_{-1}'\hat{v}_1 + \hat{\delta}'\hat{V}_{-1}'\hat{V}_2\hat{\delta} \\
&= n\left(\sum_{j=1}^{L} Z_j' Z_j \sigma_{1,j}^2 - 2\sum_{j=1}^{L} Z_j' Z_j \delta' \Sigma_{12,j} + \sum_{j=1}^{L} Z_j' Z_j \hat{\delta}' \Sigma_{2,j}\hat{\delta}\right) \\
&= n\sum_{j=1}^{L} Z_j' Z_j \left(\sigma_{1,j}^2 - 2\hat{\delta}\Sigma_{12,j} + \hat{\delta}\Sigma_{2,j}\hat{\delta}\right) \\
&= n\sum_{j=1}^{L} Z_j' Z_j \sigma_{x_{k,j}}^2
\end{aligned}
$$

The expression for variance given in section 2 of the main paper can be re-written as;

$$
\sigma_{x_{k,j}}^2 = \left(\sigma_{1,j}^2 - 2\hat{\delta}\Sigma_{12,j} + \hat{\delta}\Sigma_{2,j}^2\hat{\delta}\right)
$$

Therefore;

$$\begin{aligned}
\hat{\zeta}'\hat{\zeta} &= n\sum_{j=1}^{L} Z_j' Z_j \left(\sigma_{1,j}^2 - 2\hat{\delta}\Sigma_{12,j} + \hat{\delta}\Sigma_{2,j}\hat{\delta}\right) \\
&= n\sum_{j=1}^{L} Z_j' Z_j \sigma_{x_{k,j}}^2
\end{aligned}$$

Substituting the results obtained for the numerator and denominator into the Basmann statistic gives;

$$\begin{aligned}
B &= \frac{\sum_{j=1}^{L} Z_j' Z_j \left(\hat{\pi}_{1,j} - \hat{\Pi}_{2,j}\hat{\delta}\right)^2}{\left(n\sum_{j=1}^{L} Z_j' Z_j \sigma_{x_{1,j}}^2\right)/n} \\
&= \sum_{j=1}^{L} \frac{Z_j' Z_j \left(\hat{\pi}_{1,j} - \hat{\Pi}_{2,j}\hat{\delta}\right)^2}{Z_j' Z_j \sigma_{x_{1,j}}^2} \\
&= \sum_{j=1}^{L} \frac{\left(\hat{\pi}_{1,j} - \hat{\Pi}_{2,j}\hat{\delta}\right)^2}{\sigma_{x_{1,j}}^2} \\
&= Q_{x_k}
\end{aligned}$$

As the instruments are independent from each other. Therefore $Q_{x_k}$ is the two-sample equivalent of the Basmann statistic when the instruments are uncorrelated and consequently $F_{TS}$ is equivalent to $F_{SW}$.

### 2 Simulation results for a model with two exposures and no heterogeneity

Supplementary Table S1 gives simulation results for a model with 2 exposures and 200 SNPs as instruments. The model in this simulation is equivalent to the model used for the results given in Table 2 except for the change that here the SNPs have no pleiotropic effect on the outcome and so there is no extra heterogeneity in the model. These results show that $\hat{\beta}_Q$ and $\hat{\beta}_{Q,het}$ are equivalent when there is no heterogeneity and $\hat{\beta}_{Q,het}$ has only a very modest increase in the standard error.

Table S1: Simulation results for models with no heterogeneity: 2 exposures, 200 SNPs

| | Weak instruments | | Conditionally weak instruments | |
|---|---|---|---|---|
| | $x_1$ | $x_2$ | $x_1$ | $x_2$ |
| *Individual level estimation* | | | | |
| $\hat{\beta}_{OLS}$ | 1.09 | -0.053 | 0.77 | -0.48 |
| | (0.005) | (0.004) | (0.005) | (0.005) |
| $\hat{\beta}_{IV}$ | 0.592 | -0.328 | 0.530 | -0.321 |
| | (0.026) | (0.026) | (0.015) | (0.011) |
| $F$ | 8.81 | 8.82 | 1600.02 | 3106.6 |
| | (0.655) | (0.633) | (98.10) | (190.71) |
| $F_{IV}$ | 3.39 | 3.39 | 9.70 | 9.73 |
| | (0.356) | (0.350) | (0.92) | (0.93) |
| *Two-sample estimation with covariances* | | | | |
| $\hat{\beta}_{IVW}$ | 0.356 | -0.171 | 0.448 | -0.264 |
| | (0.041) | (0.042) | (0.027) | (0.019) |
| $\hat{\beta}_{Q}$ | 0.503 | -0.302 | 0.501 | -0.301 |
| | (0.062) | (0.062) | (0.030) | (0.022) |
| $\hat{\beta}_{Q,het}$ | 0.504 | -0.304 | 0.501 | -0.301 |
| | (0.066) | (0.063) | (0.031) | (0.022) |
| $F_{TS}$ | 3.34 | 3.35 | 9.09 | 9.12 |
| | (0.343) | (0.343) | (0.804) | (0.808) |
| *Two-sample estimation without covariances* | | | | |
| $\hat{\beta}_{IVW}$ | 0.356 | -0.171 | 0.448 | -0.264 |
| | (0.041) | (0.042) | (0.027) | (0.019) |
| $\hat{\beta}_{Q}$ | 0.514 | -0.314 | 0.741 | -0.474 |
| | (0.064) | (0.064) | (0.045) | (0.032) |
| $\hat{\beta}_{Q,het}$ | 0.517 | -0.318 | -88.15 | 63.54 |
| | (0.068) | (0.069) | 5267.32 | 3782.18 |
| $F_{TS}$ | 3.16 | 3.17 | 0.45 | 0.45 |
| | (0.332) | (0.327) | (0.049) | (0.049) |

$\beta_1 = 0.5$, $\beta_2 = -0.3$

4,000 repetitions, 20,000 observations per repetition

Covariances estimated from the correlation between $x_1$ and $x_2$

### 3 Description of the Avon longitudinal study of Parents and Children

Pregnant women resident in Avon, UK with expected dates of delivery 1st April 1991 to 31st December 1992 were invited to take part in the study. The initial number of pregnancies enrolled is 14,541 (for these at least one questionnaire has been returned or a Children in Focus clinic had been attended by 19/07/99). Of these initial pregnancies, there was a total of 14,676 foetuses, resulting in 14,062 live births and 13,988 children who were alive at 1 year of age.

When the oldest children were approximately 7 years of age, an attempt was made to bolster the initial sample with eligible cases who had failed to join the study originally. As a result, when considering variables collected from the age of seven onwards (and potentially abstracted from obstetric notes) there are data available for more than the 14,541 pregnancies mentioned above. The number of new pregnancies not in the initial sample (known as Phase I enrolment) that are currently represented on the built files and reflecting enrolment status at the age of 24 is 913 (456, 262 and 195 recruited during Phases II, III and IV respectively), resulting in an additional 913 children being enrolled. The phases of enrolment are described in more detail in the cohort profile paper and its update[7,8]. The total sample size for analyses using any data collected after the age of seven is therefore 15,454 pregnancies, resulting in 15,589 foetuses. Of these 14,901 were alive at 1 year of age.

A 10% sample of the ALSPAC cohort, known as the Children in Focus (CiF) group, attended clinics at the University of Bristol at various time intervals between 4 to 61 months of age. The CiF group were chosen at random from the last 6 months of ALSPAC births (1432 families attended at least one clinic). Excluded were those mothers who had moved out of the area or were lost to follow-up, and those partaking in another study of infant development in Avon.

The study website contains details of all the data that is available through a fully searchable data dictionary and variable search tool" and reference the following webpage: http://www.bristol.ac.uk/alspac/researchers/our-data/. Ethical approval for the study was obtained from the ALSPAC Ethics and Law Committee and the Local Research Ethics Committees. Consent for biological samples has been collected in accordance with the Human Tissue Act (2004).
